# Mass Spectrometry Imaging of Hair Identifies Daily Maraviroc Adherence in HPTN 069/ACTG A5305

**DOI:** 10.1101/2023.01.30.526384

**Authors:** Elias P. Rosen, Nicole White, William M. Gilliland, Roy R. Gerona, Monica Gandhi, K. Rivet Amico, Kenneth H. Mayer, Roy M. Gulick, Angela DM Kashuba

## Abstract

Objective measures of adherence for antiretrovirals used as pre-exposure prophylaxis (PrEP) are critical for improving preventative efficacy in both clinical trials and real-world application. Current objective adherence measures either reflect only recent behavior (eg days for plasma or urine) or cumulative behavior (eg months for dried blood spots). We measured the accumulation of the antiretroviral drug maraviroc (MVC) in hair strands by infrared matrix-assisted laser desorption electrospray ionization (IR-MALDESI) mass spectrometry imaging (MSI) to evaluate adherence behavior longitudinally at high temporal resolution. An MSI threshold for classifying daily adherence was established using clinical samples from healthy volunteers following directly observed dosing of 1 to 7 doses MVC/week. We then used the benchmarked MSI assay to classify adherence to MVC-based PrEP regimens in hair samples collected throughout the 49-week HPTN069/ACTGA5305 study. We found that only ~32% of investigated hair samples collected during the study’s active dosing period showed consistent daily PrEP adherence throughout a retrospective period of 30 days, and also found that profiles of daily individual adherence from MSI hair analysis could identify when patients were and were not taking study drug. The assessment of adherence from MSI hair strand analysis was 62% lower than adherence classified using paired plasma samples, the latter of which may be influenced by white-coat adherence. These findings demonstrate the ability of MSI hair analysis to examine daily variability of adherence behavior over a longer-term measurement and offer the potential for longitudinal comparison with risk behavior to target patient-specific adherence interventions and improve outcomes.

## Introduction

Pre-exposure prophylaxis (PrEP) with antiretroviral drugs (ARVs) is effective against HIV-1 acquisition when sufficient drug concentrations are present during periods of exposure (1–3), conditions which rely on both PrEP adherence and persistence. Based on differences in tissue pharmacokinetics, the dosing frequency required to attain protective concentrations of ARVs can vary by route of HIV exposure (4). As a result, PrEP guidelines for oral agents recommend daily use for most populations and event-driven dosing only for cisgender men engaging in anal sex. Objective measures of adherence with implementation of PrEP through clinical trials, demonstration projects, and routine clinical dissemination have shown that individual levels of adherence and persistence can be complex and often significantly lower than dosing recommendations (5–8).

Evaluating PrEP effectiveness in the context of dynamic risk behavior (9) requires information about daily changes in ARV concentrations over time which is not possible using current objective adherence measures. The period of adherence captured by pharmacologic measures varies by biological matrix based on pharmacokinetics and analysis methods. These generally fall into measurements of recent behavior (eg from plasma, saliva, or urine) or cumulative behavior (eg from intracellular metabolites in blood cells, or hair thatches) (10). Measures of recent adherence, which are susceptible to white-coat (social desirability) bias, only reflect behavior over the past few days. Cumulative measures reflect average adherence over a period of weeks to months, but cannot differentiate changes in adherence within this period. While measures can be used in combination to couple short- and long-term adherence (11, 12), these strategies are still not capable of capturing daily adherence patterns over an extended period of time reflecting patients’ actual daily use.

Hair is a unique biomatrix for adherence assessment because each strand represents a record of systemic drug concentrations incorporated into hair from blood during follicular growth and preserved as the hair grows. Sensitive analysis of hair strands by liquid chromatography-mass spectrometry (LC-MS) has been demonstrated for ARV drug concentrations, which scale proportionally with dose frequency(13) and can predict virologic success (14). LC-MS methods typically evaluate hair segments ≥ 1 cm that correspond to at least a month of growth. We have developed a new approach to measuring ARV drug exposure longitudinally along single hair strands at high spatial, and thus temporal, resolution using infrared matrix-assisted laser desorption electrospray ionization (IR-MALDESI) mass spectrometry imaging (MSI).

The HIV Prevention Trials Network (HPTN) 069/AIDS Clinical Trials Group (ACTG) A5305 study examined maraviroc (MVC) as an agent for PrEP, either alone or in combination with other antiretrovirals. Although MVC has not moved forward as a PrEP candidate, the study showed the safety and tolerability of MVC-based regimens (15, 16). In this work, we apply IR-MALDESI MSI to: 1) characterize MVC dosing behavior through benchmarking of longitudinal MVC profiles in hair following directly observed therapy (DOT) of daily and intermittent dosing in the ENLIGHTEN study; and, 2) investigate patterns of longer-term adherence in HPTN069/ACTGA5305 study samples, comparing these measures to commonly used adherence assessments. Using MVC as proof-of-concept, we demonstrate the capability of MSI hair analysis to examine daily ARV adherence patterns over one month via a single assay.

## Results

### IR-MALDESI MSI benchmarking of MVC in hair strands: The ENLIGHTEN Study

The ENLIGHTEN directly-observed-therapy (DOT) study provided MVC to HIV-negative volunteers in different dosing patterns. In our assessment of MVC disposition by IR-MALDESI MSI through the ENLIGHTEN study, we found that the quantitative patterns of MVC detectable along hair strands were well aligned with known dosing information, which ranged from daily (7x/week) to interrupted therapy (0, 1 or 3x/week). Regions of MVC accumulation associated with hair growth during an interrupted dosing period (3x/week, 1x/week, or 0x/week) can be seen in Fig. 1A, along with higher MVC response on the right-hand, distal portion of the hair strands corresponding to growth during an earlier period of daily (7x/week) dosing. The 7-day washout interval between daily and differentiated dosing is also apparent, particularly between daily and 3x/week dosing shown at the top of Fig. 1A. A time series illustrating the movement of drug distally in hair strands collected throughout the transition between daily and intermediate dosing periods for one subject is shown in Fig. S1.

**Fig. 1.**
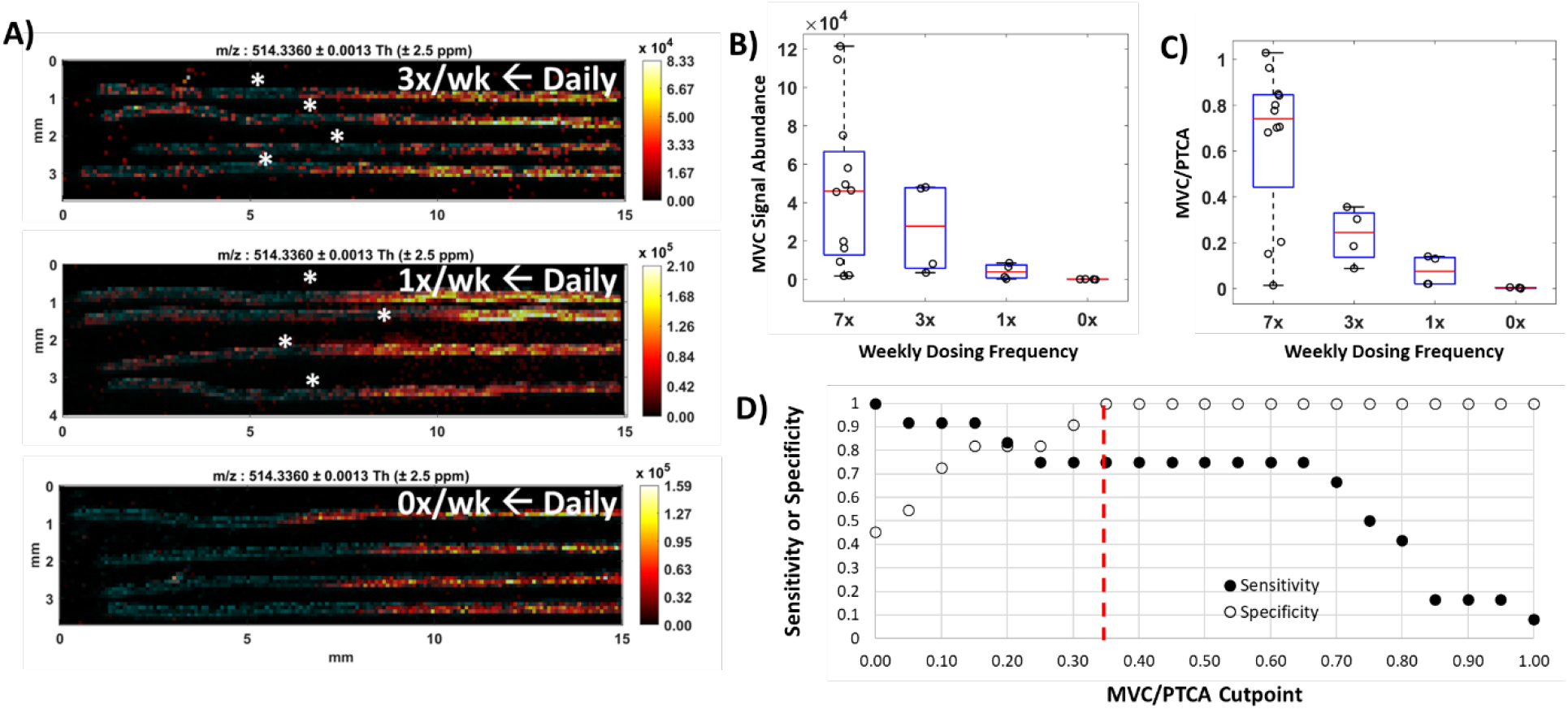
Benchmarking MVC in hair strands with MSI. (A) Representative IR-MALDESI MSI MVC ion maps showing drug accumulation associated with daily dosing and each intermittent dosing group (from top: 3x/week, 1x/week, and 0x/week, respectively) in hair strands oriented with time of growth increasing distally from left to right. MVC was measured over the proximal 15 mm, an estimated 1.5 months of growth, of samples collected at the end of the intermittent dosing period to evaluate dose-response of MVC accumulation in hair. MVC signal abundance (m/z 514.3360) is represented by a color scale increasing in concentration from regions of dark red/black to regions of orange/yellow. Cholesterol present in the hair strands (m/z 369.3516, shown in blue) is overlaid to clearly show the shape, orientation, and length of the individual strands. Apparent regions of each strand associated with a 7-day washout between dosing periods are denoted by a white asterisk. Hair was collected by clipping close to the scalp. (B) Mean IR-MALDESI MVC signal abundance associated with each ENLIGHTEN dosing group evaluated from composite longitudinal hair profiles. (C) PTCA-normalized MVC signal abundance associated with each dosing group. (D) ROC sensitivity and specificity of daily MVC adherence binary classification based on adherence cutpoint. The selected cutpoint value MVC/PTCA=0.35 is demarcated by a red dashed line.

While within-individual longitudinal profiles showed MVC response scaling with dosing frequency, this observation did not hold across the whole cohort. Interquartile ranges of mean MVC signal abundance from daily and intermittent dosing regions could not be differentiated (Fig. 1B). Volunteers had a range of hair colors (table S1) and to account for between subject variability in MVC accumulation this caused, we normalized the raw MVC signal abundance by a melanin biomarker (pyrrole-2,3,5-tricarboxylic acid, PTCA) measured in the same hair strands. Mean MVC/PTCA values (Fig. 1C) had interquartile ranges for each dosing frequency that could be differentiated [MVC/PTCA, Daily: 0.745(0.440-0.845); 3x: 0.245 (0.140-0.330); 1x: 0.075(0.020-0.135); 0x: 0.002 (0.000-0.005)].

Selection of a threshold value for PTCA-normalized MVC signal abundance for binary classification of adherence (adherent to regimen vs. not adherent to regimen) was determined from a receiver operating characteristic curve. Fig. 1D shows the relationship between MVC/PTCA threshold values and the true positive rate (sensitivity) and true negative rate (specificity) of binary classification. We selected a threshold value of MVC/PTCA=0.35 (specificity: 100%; sensitivity: 75%) to differentiate 3 or fewer doses per week from more frequent dosing behavior, prioritizing high specificity to minimize incorrectly labeling a patient as non-adherent to medication which could damage patient motivation (17).

### Assessing MVC PrEP adherence patterns in HPTN069/ACTGA5305

Cumulative MVC concentrations in hair strands collected from across the three HPTN069/ACTGA5305 hair sampling timepoints of a pharmacologic substudy (18) (on drug: Week 24 and 48; follow-up: Week 49; table 1) were found to be strongly correlated between measurements by MSI and LC-quadrupole time-of-flight mass spectrometry (LC-QTOF/MS) (Fig. 2A; Spearman’s rho, r = 0.78, P<0.0001). MVC detectability (MSI and LC-QTOF/MS: 26/32 measurable samples), medians (MSI: 0.394 ng/mg, LC-QTOF/MS: 0.361 ng/mg) and ranges (MSI: 0.120-2.03 ng/mg, LC-QTOF/MS: 0.035-1.53 ng/mg) were similar across methods over matched hair segment lengths. We found no difference in MVC concentration among regimen arms that included MVC (Fig. 2B; P>0.14) or between sexes (Fig. 2C; P=0.94).

**Table 1.** Summary of investigated HPTN069/ACTGA5305 hair samples.

| Dosing Arm | Study Week |  |  | Longitudinal |
| --- | --- | --- | --- | --- |
|  | *On drug* |  | *Follow-up* | Samples from |
|  | 24 | 48 | 49 | Individual Patients |
| *Maraviroc* |  |  |  |  |
| Male | 2 | 2 | 2 | 2 |
| Female | 2 | 1 | 0 | 0 |
| *Maraviroc+Emtricitabine* |  |  |  |  |
| Male | 4 | 2 | 0 | 2 |
| Female | 3 | 2 | 1 | 1 |
| *Maraviroc+Tenofovir* |  |  |  |  |
| *Disoproxil Fumarate* |  |  |  |  |
| Male | 4 | 1 | 2 | 1 |
| Female | 1 | 1 | 2 | 1 |
| Total | 16 | 9 | 7 | 7 |

**Fig. 2.**
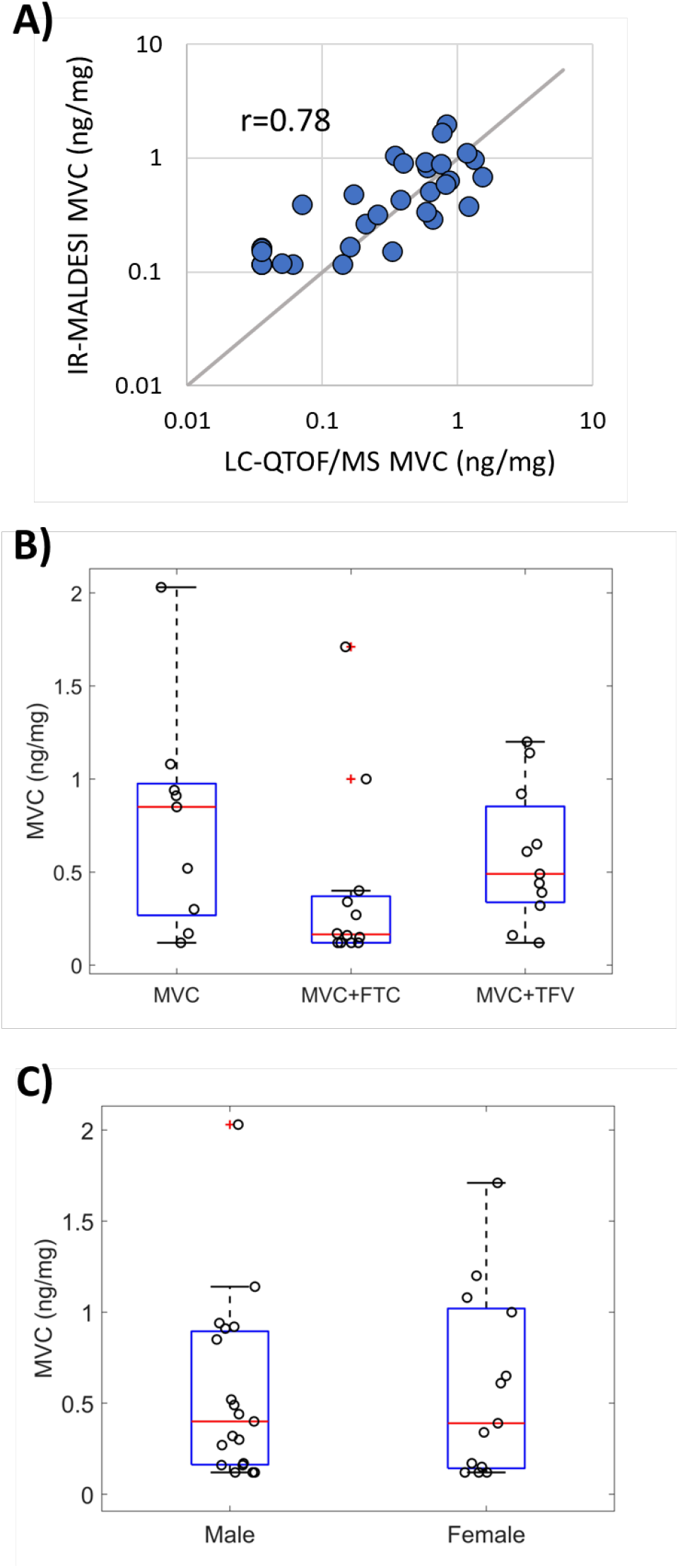
Cumulative MVC accumulation in hair strands during HPTN069/ACTGA5305. (A) Cumulative MVC concentration evaluated in the proximal 1 cm of hair strands by LC-QTOF/MS and MSI. Gray 1:1 line provided for comparison. (B) HPTN069/ACTGA5305 hair substudy MVC concentrations in each dosing arm. (C) HPTN069/ACTGA5305 hair substudy MVC concentrations by sex.

Using the MVC/PTCA threshold, classification of adherence behavior among the participants (table S2) over the prior 30 days fell into three groups: drug response consistent with no days of adherence (n=12 blue points on left side of Fig. 3A; “no days”); drug response consistent with some days of adherence (n=9 blue points across center of Fig. 3A; “some days”); and, drug response consistent with all days of adherence (n=11 blue points on right side of Fig. 3A; “all days”). As with benchmarking, we see in Fig. 3A that such classification groupings are more ambiguous by MVC concentration alone: a cumulative hair concentration of 0.61 ng/mg, for example, was measured in samples classified separately as having 0/30 and 30/30 days of adherence, respectively. The cumulative MVC concentrations among samples classified as having some days of adherence are not significantly different from those with all days of adherence (Fig. 3B; P=0.61). Conversely, normalization of MVC by PTCA results in MVC/PTCA distributions that are distinguishable across adherence groups (Fig. 3C; no days:some days, P =0.028; no days:all days, P < 0.001; and, some days: all days, P= 0.067). The analysis of daily adherence behavior made possible through IR-MALDESI MSI also reveals variation in the individual patterns of normalized drug responses – including numbers of consecutive days of non-adherence – over the 30-day observation period among the group of samples with “some days” of adherence (Fig. 3D). Accompanying bar graphs for samples categorized as “no days” and “all days” of adherence are provided in Fig. S2 and S3, respectively.

**Fig. 3.**
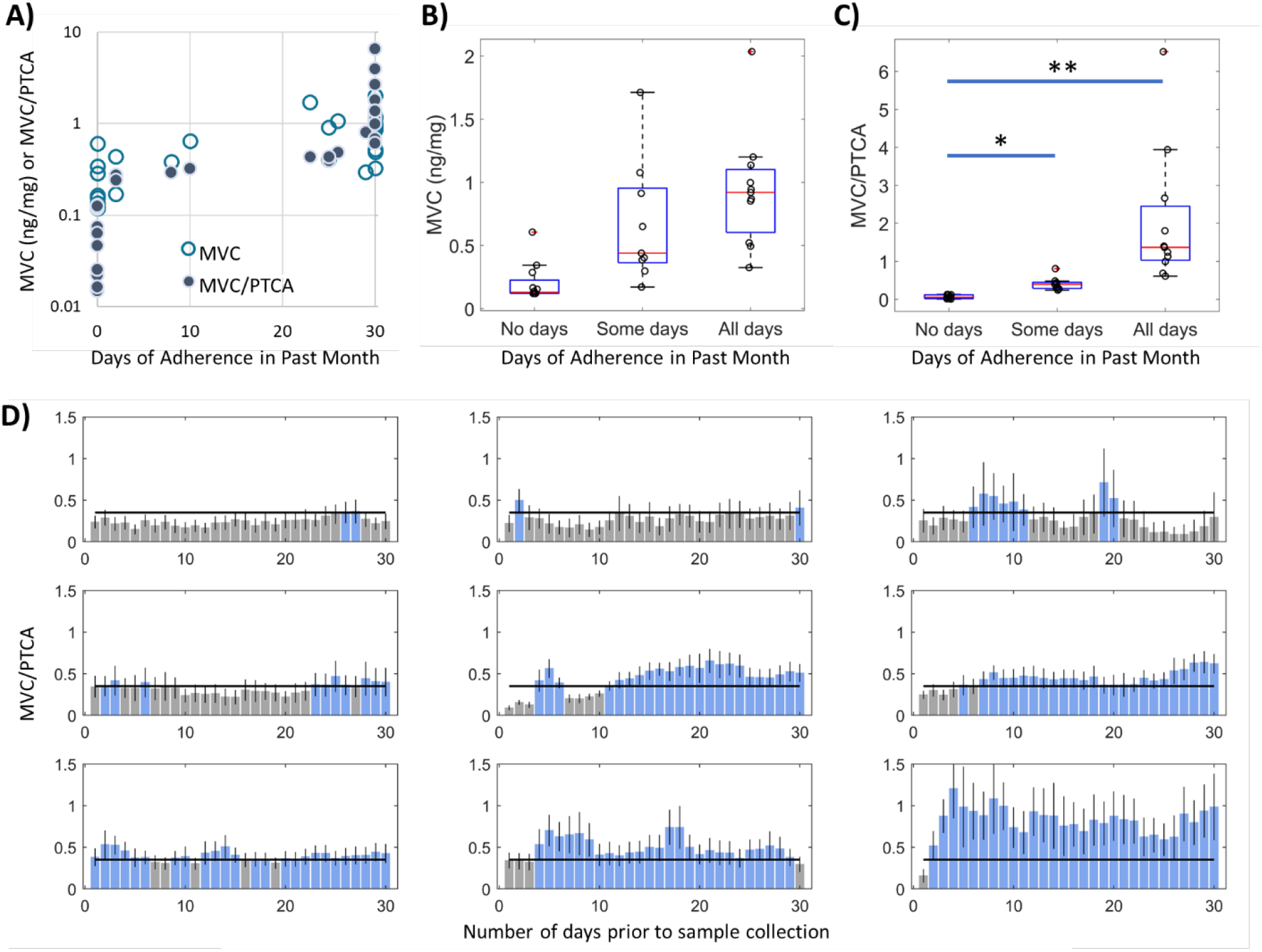
Adherence classification of HPTN069/ACTGA5305 hair strands. (A) MVC concentration or MVC/PTCA normalized response in hair strands relative to the number of days classified as reflecting adherence within the past 30 days prior to hair sample collection. (B) MVC concentration of hair strands within groups of adherence behavior associated with no, some, or all days classified as adherent. (C) MVC/PTCA response in hair strands within groups of adherence behavior associated with no, some, or all days classified as adherent. (D) Bar graphs of adherence measured by MSI within samples where “some days” of adherence was determined. Blue bars reflect days in which MVC response exceeded the adherence cutpoint and gray bars reflect days where MVC response did not exceed the adherence cutpoint.

Cumulative concentrations of MVC in hair over the prior month had poor correlation with plasma concentrations of MVC, which have an elimination half-life of 16 hours, in matched samples when participants were on study drugs (weeks 24 and 48) and at follow-up (week 49) (Fig. 4A; r = −0.07, P=0.72). Correlation between drug concentrations in hair assessed via LC-QTOF/MS to plasma MVC measures was similarly low (r = −0.03, P=0.87). Correlation between these measures improved when comparing IR-MALDESI MSI and plasma in weeks that participants were on drug (i.e., weeks 24 and 48) to account for the rapid MVC clearance from plasma after PrEP discontinuation, which occurred at week 49 (r =0.40, P=0.05). Further comparison of hair and plasma results at 24 and 48 weeks reveals disagreement in binary classification of adherence (Fig. 4B), with 84% of samples (21/25) classified as adherent by the plasma MVC threshold of 4.6 ng/ml (18) and only 32% of samples (8/25) classified as adherent according to IR-MALDESI. Agreement between the two classification measures occurred in only 32% of samples, and 88% of discordance occurred in samples classified as adherent based on plasma and non-adherent based on hair analysis (McNemar test: χ^2^=9.94, P=0.0016). Comparison of whole-cohort MVC concentrations in hair at week 24 vs. week 48 (Fig. 4C, left) shows no statistically significant difference between time points (P=0.73). In the seven participants for whom data were available at both 24 and 48 weeks, one patient had an apparent increase in MVC concentration between week 24 and 48, two had low concentrations at both time points, and four had a decrease in hair concentrations. As with Fig. 3C, we find that normalization to account for different melanin levels reveals different patterns from those in unadjusted analyses (cf. right panel vs. left panel of Fig. 4C): three participants appear to have similar levels of adherence at 24 and 48 weeks, two have consistent non-adherence, and two have decreasing adherence. For these seven participants, agreement between plasma and MSI hair adherence classification at 24 and 48 weeks varied from total discordance to total concordance (Fig. 4D).

**Fig. 4.**
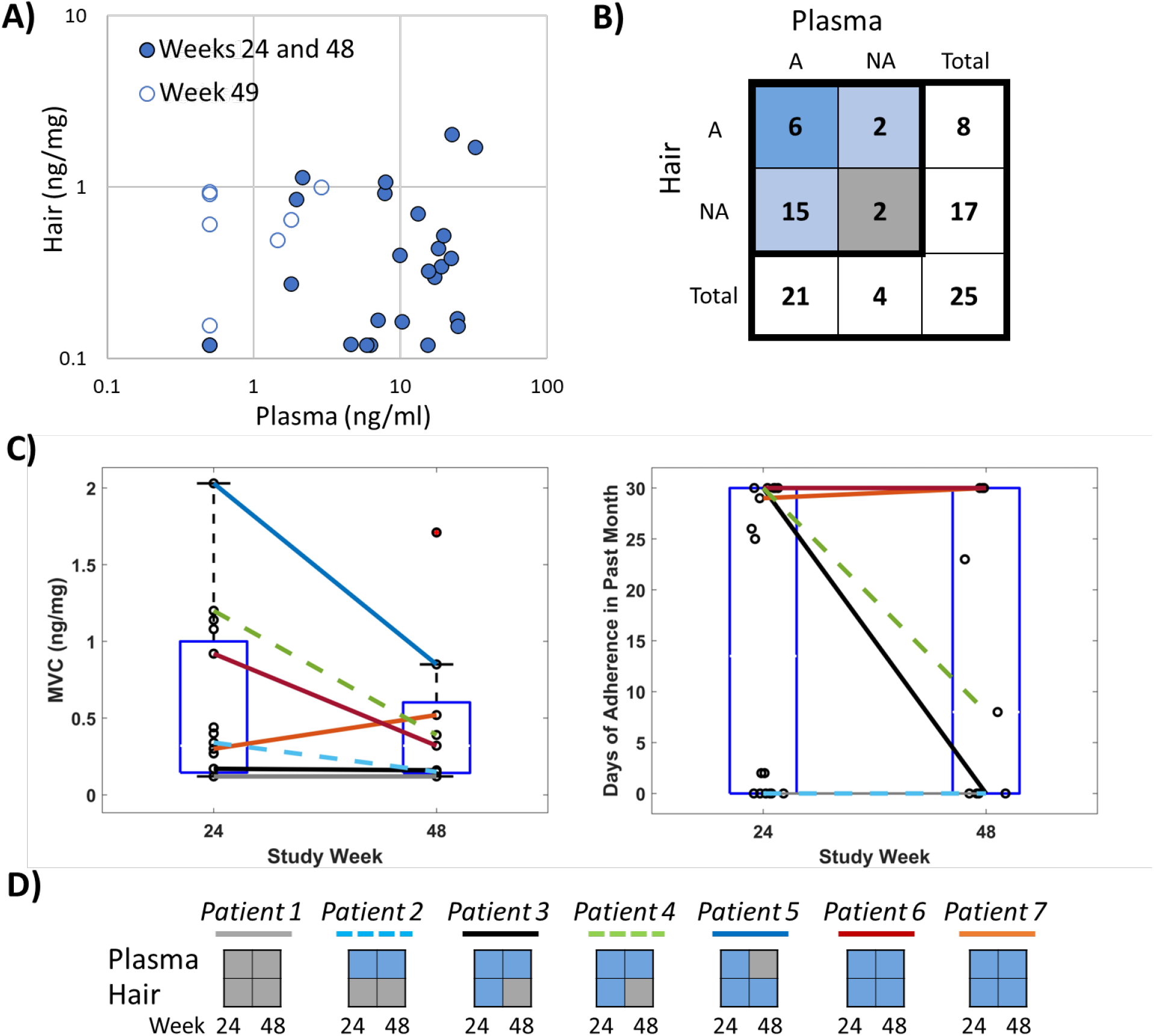
Comparison of long-term and short-term objective adherence measures from HPTN069/ACTGA5305. (A) Comparison of plasma concentration and hair MSI concentration. Samples collected at week 24 and 48 are denoted by a solid circle and samples collected at week 49 have an open circle. (B) Contingency table of adherence classification by hair and plasma where *A* denotes the number of samples classified as adherent and *NA* denotes the number of samples classified as non-adherent. Table shading reflects agreement between measurements where blue corresponds to matched classification of adherence, light blue corresponds to discordance in adherence classification, and gray corresponds to matched classification on non-adherence. (C) Comparison of hair MSI MVC concentration (left) and adherence assessment (right) for on-drug samples collected from individuals at both weeks 24 and 48. Patients with samples at both weeks (n=7) are denoted by a colored line that is solid (male) or dashed (female). (D) Comparison in longitudinal adherence classification (blue, adherent; gray, non-adherent) between hair and plasma for 7 patients with both types of samples available at each visit.

Poor agreement was also found between frequency of pill openings within the prior month measured by Wisepill electronic monitoring and pharmacologic measures of adherence in 15 samples with matched records (Fig. S4). Spearman’s rho relating pill openings to hair MVC concentrations (r =0.12, P=0.68) and plasma (r =−0.07, P=0.81) suggest low correlation between paired values.

## Discussion

IR-MALDESI MSI was able to evaluate MVC longitudinally in single hair strands at high spatial resolution and classify short-term changes in adherence behavior over the longer-term drug dosing record provided by hair. Accounting for differences in the accumulation of MVC in hair based on its melanin content was necessary to unambiguously classify adherence in the ENLIGHTEN study. Correlation between melanin and concentration of drug in hair has been well documented in forensic toxicology analysis of illicit drugs (19, 20), where the effect is stronger for basic compounds. Our prior work showed that a biomarker of melanin, PTCA, was more strongly correlated with MVC accumulation in hair than more acidic antiretrovirals such as emtricitabine and dolutegravir and suggested that the binding of MVC to melanin may limit its removal from hair strands after chemical hair treatments (21). Binary classification of adherence based on MVC/PTCA normalization was selected to maintain assay applicability across varied hair colors and types, and we prioritized assay specificity in establishing a MVC/PTCA threshold cutpoint differentiating 3 or fewer doses/week from more frequent dosing to avoid misclassification of non-adherence (17). As a result, adherence classification based on the selected cutpoint can be considered conservative for differentiating daily and intermittent dosing. Since target concentrations of MVC for PrEP have not been defined, this cutpoint only reflects adherence and not efficacy.

Longer-term adherence in HPTN069/ACTGA5305 hair strands provided a considerably different perspective than adherence assessed in plasma samples collected at the same timepoints. Daily adherence classification by IR-MALDESI MSI indicated that less than one-third of hair samples reflected consistent, sustained adherence throughout the prior 30 days. Matched plasma MVC concentrations, classified by an adherence threshold defined conservatively relative to benchmarked TFV and FTC plasma adherence thresholds (18), indicated much higher recent adherence (84%). These samples may not represent an overlapping period of drug exposure because the proximal end of cut hair may correspond to growth occurring more than a week before collection (22), so comparisons have been interpreted solely as differences in short-term and longer-term adherence behavior. Higher adherence classification in plasma samples likely arises from changes in adherence behavior just prior to a clinic visit such as white-coat adherence (23) or participants simply being sensitized to the study and related procedures around study visits. The discordance between these measures underscores the need for monitoring both short and longer-term adherence in evaluations of PrEP efficacy.

Findings of longer-term adherence captured in hair were consistent with an additional pharmacologic assessment in the HPTN069/ACTGA5305 tissue sub-study. A comparison of the interquartile range of rectal concentrations of PrEP antiretrovirals (MVC, TFV, and FTC) (18) with pharmacokinetic studies of rectal drug concentrations ranging from single dose to steady-state dosing (24–26) (table S4) indicates that median tissue concentrations from individuals sampled across each dosing arm of HPTN069/ACTGA5305 fell below median values expected from daily dosing and may be more consistent with 4 or fewer doses per week. Levels of adherence suggested by these tissue concentrations are similar to our IR-MALDESI findings but not the high short-term adherence indicated by plasma.

Hair strand MSI analyses indicated heterogeneous periods of active dosing over 30 days with periodic PrEP engagement quantifiable in all but 5 samples. This approach offers a unique ability to evaluate both short-term and longer-term changes that are not captured by Wisepill data, which were not well-correlated with any other pharmacologic measures of adherence in HPTN069/ACTGA5305. It is important to note that none of the hair samples investigated here came from the 6 individuals who seroconverted during HPTN069/ACTGA5305 (15). While the number of individuals for whom we were able to evaluate samples collected from both timepoints during active PrEP dosing in the HPTN069/ACTGA5305 study was limited, we found significant differences in adherence patterns that remained either high or low, or decreased over the course of the study. Although reasons behind non-adherence are varied, some patterns of use identified by hair MSI may have been a prevention-effective strategy (9), whereby study subjects took PrEP during periods of potential exposure to HIV, or perceived risk of HIV. A recent investigation of sexual behavior among MSM within the HPTN069/ACTGA5305 cohort indicated that participants reporting condom-less sex had higher rates of plasma drug concentrations classified as adherent (27). Assessing adherence in the context of risk behavior may provide an important mechanism to support and sustain adherence and persistence of PrEP use.

Our study has several limitations. The sample size of our benchmarking study was small, which allowed us to identify a threshold differentiating daily dosing from 3 or fewer doses per week but further studies will be needed to discriminate dosing frequency more granularly. Collection of hair strands by cutting precluded interrogation of an individual’s most recent drug-taking behavior, and we recommend plucking 5 strands when assessment of the most recent week of dosing is essential. Finally, the availability of HPTN069/ACTGA5305 hair samples limited the number of individuals who participated in the pharmacokinetic sub-study that we could investigate.

We have shown the unique capabilities of IR-MALDESI MSI for evaluating daily antiretroviral adherence throughout the record of drug accumulation preserved in hair strands. The approach is sensitive to MVC as well as a range of ARVs and other small molecules (28) making it highly adaptable for monitoring multidrug regimens, including those containing FTC as we have demonstrated previously (29). IR-MALDESI MSI offers a new approach for measuring adherence patterns that provides a temporal overview of dosing that can be used in research and could be an important addition to adherence monitoring and intervention.

## Methods

### Study Design

Benchmarking of MVC in hair strands was performed as part of the ENLIGHTEN Study (NCT03218592). Consenting HIV-uninfected healthy volunteers (n=12, table S1) were administered MVC 300mg by directly observed therapy. All study volunteers participated in a 28-day period of daily dosing after which they were randomized (n=4) for a subsequent 28-day period to one of three differentiated dosing frequencies: 0 doses/week, 1 dose/week, or 3 doses/week. An interval of 7 days separated each dosing period. Hair was collected by cutting approximately 10 hair strands from the occipital region close to the scalp using scissors, adhered to aluminum foil at their distal end to preserve orientation, and stored with a desiccant gel pack at 4°C until analysis. MSI response to MVC accumulation in ENLIGHTEN hair strands in daily and intermittent dosing periods was evaluated using samples collected at the end of each phase.

Characterization of PrEP adherence by IR-MALDESI MSI was performed through HPTN069/ACTGA5305 (NCT01505114). This was a 48-week placebo-controlled study in at-risk MSM and women of the safety and tolerability of candidate HIV PrEP regimens including MVC alone or in combination with either tenofovir disoproxil fumarate (TDF) or emtricitabine (FTC) in comparison to TDF+FTC, conducted from 2012-2015 (15, 16). Adherence measurements were undertaken for all participants (electronic drug monitoring using a pillbox (Wisepill) containing the 3 ARVs, drug level monitoring from blood stored at every visit), with additional sampling of tissues, plasma, and hair conducted through a nested pharmacologic substudy (18). Approximately 200 strands of hair were collected from sub-study participants at three time points (on drug: Week 24, Week 48; follow-up: Week 49). Hair storage followed the same protocol as ENLIGHTEN. IR-MALDSI MSI analysis was performed on MVC-based regimen samples (MVC, MVC+TDF, MVC+FTC) for which hair was not consumed during LC-MS analysis. As summarized in Table 1, this corresponded to a total of 32 samples collected across the study from 19 participants (10 male, 9 female) whose demographic information is included in table S2.

### Hair Analysis

#### IR-MALDESI MSI

Hair strands (n=4) were oriented horizontally and adhered to glass microscope slides with proximal strand ends positioned to the left for analysis by IR-MALDESI MSI (28, 30). Longitudinal analysis of MVC was performed using a two-step laser desorption and electrospray ionization process has been detailed elsewhere (21, 30) and will be described briefly. Prepared sample slides were positioned on a temperature-controlled stage in the IR-MALDESI MSI source enclosure before being cooled to −9°C under dry nitrogen gas flow to reduce humidity. Following temperature stabilization, the nitrogen flow was interrupted and the MSI source was opened to the ambient atmosphere to grow a thin layer of ice on the sample surface. Following ice growth, the source was closed and nitrogen was used to maintain a relative humidity of ~14% throughout the experiment to preserve ice thickness. The ice layer promoted sample desorption from single IR laser pulses (*λ*=2940 nm, IR Opolette, Opotek, Carlsbad, CA). Volatilized material expanding upward from the sample intersected an orthogonal electrospray plume to create analyte ions, which were sampled into an orbitrap mass spectrometer (ThermoFisher Q Exactive Plus, Bremen, Germany) for analysis. A list of targeted analytes is shown in table S4. For analysis of positive ions, the mass spectrometer was operated in positive polarity full scan mode (m/z 200 to 800; resolving power: 140,000 at m/z 200; s-lens RF level: 50, mass accuracy: <1 ppm). For analysis of negative ions, the mass spectrometer was operated in full scan mode with negative polarity (*m/z* 190 – 760; resolving power: 140,000 at *m/z* 200; s-lens RF level: 50, mass accuracy: <1 ppm). Analysis was performed with a step-size of 100 μm between sampling locations, corresponding to approximately 7-8 hours of growth based on the average growth rate (~ 1cm/month) in the occipital region (22). A summary of acquisition parameters and a list of targeted analytes is shown in table S3. Separate regions of interest were interrogated by IR-MALDESI MSI for analysis of MVC and the melanin biomarker PTCA (fig. S5). MVC was evaluated proximally, corresponding to the most recent growth of hair prior to sampling, and PTCA was evaluated distally by submerging this end of the slide in a solution of 1 M ammonium hydroxide in 45/45/10 methanol/water/hydrogen peroxide (v/v/v) for 10 min (21).

Calibration of IR-MALDESI response to MVC in hair strands was performed using standards prepared from blank (drug-free) hair matrix by incubation in drug, covering the range 0.145-2.99 ng/mg hair. Standards were prepared by transferring drug-free hair (approximately 10 mg) into a vial containing 20 mL of analyte and solvent (50:50 Methanol:Water), incubating for approximately 24 hours in a reciprocal shaking bath before hair was rinsed with fresh solvent and stored at −20°C. One level of standards (0.299 ng/mg) was reserved for use as a positive control in all assessments of clinical samples. A representative image showing the MVC response from a calibration and a composite calibration curve from n=5 calibrations conducted during experimental work is shown in fig. S6.

Data were processed using MSiReader and custom MATLAB software (Mathworks, Inc., Natick, MA) (30). The MVC response from three neighboring sampling locations along a composite longitudinal profile of sampled hair strands was binned to evaluate daily accumulation of drug in hair throughout the entire period of assessment (benchmarking: 1.5 cm, 1.5 months; HPTN069/ACTGA5305: 1 cm, 1 month). Normalized MVC/PTCA profiles were compared to the adherence threshold for daily adherence classification. Cumulative concentrations of MVC were determined by averaging MSI signal abundance over a segment length matched to LC-MS analysis.

#### LC-QTOF/MS

LC-MS analysis of MVC in HPTN069/ACTGA5305 hair samples was conducted in the UCSF TB Hair Analysis Laboratory using an Agilent Liquid Chromatograph 1260 (Agilent Technologies, Sta Clara, CA) attached to an Agilent Quadrupole Time-of-Flight Mass Spectrometer 6550 (31). Hair strands (proximal 1 cm segments, 2 mg) were pulverized using an Omni Bead Ruptor homogenizer (OMNI International, NW Kennesaw, GA, USA). Pulverized hair was extracted with 0.5 mL methanol followed by a two-hour mixing in a water bath shaker maintained at 37°C; the resulting extract was evaporated before reconstitution to 0.2 mL 10% acetonitrile in water with 0.1% formic acid. The sample extract (5 μL) was injected into the Agilent Liquid Chromatograph 1260 (Agilent Technologies, Sta Clara, CA) attached to an Agilent Quadrupole Time-of-Flight Mass Spectrometer 6550. Analytes in the sample extract were separated by gradient elution on an Agilent Poroshell 120, EC-C18 column (2.1 × 100 mm, 2.7 μm particle size) using water with 0.05% formic acid and 5mM ammonium formate as mobile phase A (MPA) and acetonitrile with 0.05% formic acid as mobile phase B (MPB). The gradient used for analyte separation consisted of 5% MPB at 0–0.5 min, gradient to 30% MPB from 0.5 to 1.5 min, gradient to 70% MPB from 1.5 to 4.5 min, gradient to 100% MPB at 4.5-7.5 min, and 100% MPB at 7.5–10 min; a post-wash at 5% MPB followed each run for 4 min. Ionization of MVC in the mass spectrometer was achieved using electrospray ionization (ESI) in positive polarity, and data acquisition was performed in the auto- MS/MS mode. Detection of the analyte was done by accurate mass match within 10 parts per million, retention time match within 0.1 min, target score (indicator of isotopic pattern match) of at least 70 and an MS/MS spectral match score of at least 70.

Quantification of MVC was done by isotope dilution method using MVC-d6 as internal standard. MVC drug levels were normalized by weight. The limit of detection was 0.05 ng/mg while the lower limit of quantification was 0.2 ng/mg. Procedural quality control materials and procedural blank were run along with the calibration curve at the start, middle, and end of each run. Two quality control materials were used at low and high concentrations. To accept the results of a batch run, QC materials measurements must be within 15% of their target values.

### Statistical Analyses

The Wilcoxon rank-sum test was used to compare two experiment groups. Spearman’s rank order correlation was used to assess the relationship between two analytical measures of matched samples. A Kruskal-Wallis one-way analysis of variance (ANOVA) test followed by Dunn-Sidak p-value corrections for multiple comparisons was performed between three experiment groups. Statistical significance by these methods was obtained by using Matlab. McNemar’s test of paired nominal data was used to compare binary adherence classifiers, obtained from R.

All IR-MALDESI MSI data are available online through the METASPACE database.

## Acknowledgements

This work was supported in part by the National Institutes of Allergy and Infectious Diseases and National Institutes of Mental Health (P30 AI50410, R01 AI122319), AIDS Clinical Trial Group UM1 AI 068636, and HIV Prevention Trial Network UM1-AI068619. We gratefully acknowledge the participants and members of the HPTN 069/ACTG A5305 study team. We would especially like to thank Krista Yuhas for providing demographic and electronic monitoring data.

## Supporting Information

Figure S1. Time-series of MVC accumulation in hair strands following the transition from daily to differentiated dosing.

Figure S2. Bar graphs of daily adherence classification measured by MSI within samples where “no days” of adherence was determined.

Figure S3. Bar graphs of daily adherence classification measured by MSI within samples where “all days” of adherence was determined.

Figure S4. Correlation between Wisepill frequency of pill openings and MVC concentrations in hair or plasma.

Figure S5. MSI sample analysis approach.

Figure S6. MSI MVC calibration with incubated hair standards.

Table S1. Clinical information on healthy individuals administered maraviroc in the ENLIGHTEN study.

Table S2. Clinical information on participants of HPTN069 providing hair samples.

Table S3. IR-MALDESI MSI analytes targeted in hair strand analysis and acquisition parameters Table S4. Colorectal tissue concentrations of ARVs in HPTN069 compared to PK studies.

## References

1. Baeten JM, Donnell D, Ndase P, Mugo NR, Campbell JD, Wangisi J, Tappero JW, Bukusi EA, Cohen CR, Katabira E, Ronald A, Tumwesigye E, Were E, Fife KH, Kiarie J, Farquhar C, John-Stewart G, Kakia A, Odoyo J, Mucunguzi A, Nakku-Joloba E, Twesigye R, Ngure K, Apaka C, Tamooh H, Gabona F, Mujugira A, Panteleeff D, Thomas KK, Kidoguchi L, Krows M, Revall J, Morrison S, Haugen H, Emmanuel-Ogier M, Ondrejcek L, Coombs RW, Frenkel L, Hendrix C, Bumpus NN, Bangsberg D, Haberer JE, Stevens WS, Lingappa JR, Celum C. 2012. Antiretroviral Prophylaxis for HIV Prevention in Heterosexual Men and Women. New England Journal of Medicine 367:399–410.

2. Grant RM, Lama JR, Anderson PL, McMahan V, Liu AY, Vargas L, Goicochea P, Casapía M, Guanira-Carranza JV, Ramirez-Cardich ME, Montoya-Herrera O, Fernández T, Veloso VG, Buchbinder SP, Chariyalertsak S, Schechter M, Bekker L-G, Mayer KH, Kallás EG, Amico KR, Mulligan K, Bushman LR, Hance RJ, Ganoza C, Defechereux P, Postle B, Wang F, McConnell JJ, Zheng J-H, Lee J, Rooney JF, Jaffe HS, Martinez AI, Burns DN, Glidden DV. 2010. Preexposure Chemoprophylaxis for HIV Prevention in Men Who Have Sex with Men. New England Journal of Medicine 363:2587–2599.

3. Choopanya K, Martin M, Suntharasamai P. 2013. Lancet 381:2083.

4. Cottrell ML, Yang KH, Prince HMA, Sykes C, White N, Malone S, Dellon ES, Madanick RD, Shaheen NJ, Hudgens MG, Wulff J, Patterson KB, Nelson JAE, Kashuba ADM. 2016. A Translational Pharmacology Approach to Predicting Outcomes of Preexposure Prophylaxis Against HIV in Men and Women Using Tenofovir Disoproxil Fumarate With or Without Emtricitabine. Journal of Infectious Diseases 214:55–64.

5. Marrazzo JM, Ramjee G, Richardson BA, Gomez K, Mgodi N, Nair G, Palanee T, Nakabiito C, van der Straten A, Noguchi L, Hendrix CW, Dai JY, Ganesh S, Mkhize B, Taljaard M, Parikh UM, Piper J, Mâsse B, Grossman C, Rooney J, Schwartz JL, Watts H, Marzinke MA, Hillier SL, McGowan IM, Chirenje ZM. 2015. Tenofovir-Based Preexposure Prophylaxis for HIV Infection among African Women. New England Journal of Medicine 372:509–518.

6. Van Damme L, Corneli A, Ahmed K, Agot K, Lombaard J, Kapiga S, Malahleha M, Owino F, Manongi R, Onyango J, Temu L, Monedi MC, Mak’Oketch P, Makanda M, Reblin I, Makatu SE, Saylor L, Kiernan H, Kirkendale S, Wong C, Grant R, Kashuba A, Nanda K, Mandala J, Fransen K, Deese J, Crucitti T, Mastro TD, Taylor D. 2012. Preexposure Prophylaxis for HIV Infection among African Women. New England Journal of Medicine 367:411–422.

7. van der Straten A, Van Damme L, Haberer JE, Bangsberg DR. 2012. Unraveling the divergent results of pre-exposure prophylaxis trials for HIV prevention. Aids 26:F13–F19.

8. Zhang J, Li C, Xu J, Hu Z, Rutstein SE, Tucker JD, Ong JJ, Jiang Y, Geng W, Wright ST, Cohen MS, Shang H, Tang W. 2022. Discontinuation, suboptimal adherence, and reinitiation of oral HIV pre-exposure prophylaxis: a global systematic review and meta-analysis. The Lancet HIV 9:e254–e268.

9. Haberer JE, Bangsberg DR, Baeten JM, Curran K, Koechlin F, Amico KR, Anderson P, Mugo N, Venter F, Goicochea P, Caceres C, O’Reilly K. 2015. Defining success with HIV pre-exposure prophylaxis: A prevention-effective adherence paradigm. AIDS 29:1277–1285.

10. Spinelli MA, Haberer J, Chai PR, Castillo-Mancilla J, Anderson PL. Approaches to Objectively Measure Antiretroviral Medication Adherence and Drive Adherence Interventions. Current HIV/AIDS reports 17:301–314.

11. Castillo-Mancilla JR, Zheng JH, Rower JE, Meditz A, Gardner EM, Predhomme J, Fernandez C, Langness J, Kiser JJ, Bushman LR, Anderson PL. 2013. Tenofovir, emtricitabine, and tenofovir diphosphate in dried blood spots for determining recent and cumulative drug exposure. AIDS Research and Human Retroviruses 29:384–390.

12. Velloza J, Bacchetti P, Hendrix CW, Murnane P, Hughes JP, Li M, Curlin ME, Holtz TH, Mannheimer S, Marzinke MA, Amico KR, Liu A, Piwowar-Manning E, Eshleman SH, Dye BJ, Gandhi M, Grant RM, Team HAS. 2019. Short- and Long-Term Pharmacologic Measures of HIV Pre-exposure Prophylaxis Use Among High-Risk Men Who Have Sex With Men in HPTN 067/ADAPT. Jaids-Journal of Acquired Immune Deficiency Syndromes 82:149–158.

13. Liu AY, Yang QY, Huang Y, Bacchetti P, Anderson PL, Jin CS, Goggin K, Stojanovski K, Grant R, Buchbinder SP, Greenblatt RM, Gandhi M. 2014. Strong Relationship between Oral Dose and Tenofovir Hair Levels in a Randomized Trial: Hair as a Potential Adherence Measure for Pre-Exposure Prophylaxis (PrEP). Plos One 9.

14. Gandhi M, Ameli N, Bacchetti P, Anastos K, Gange SJ, Minkoff H, Young M, Milam J, Cohen MH, Sharp GB, Huang Y, Greenblatt RM. 2011. Atazanavir Concentration in Hair Is the Strongest Predictor of Outcomes on Antiretroviral Therapy. Clinical Infectious Diseases 52:1267–1275.

15. Gulick RM, Wilkin TJ, Chen YQ, Landovitz RJ, Amico KR, Young AM, Richardson P, Marzinke MA, Hendrix CW, Eshleman SH, McGowan I, Cottle LM, Andrade A, Marcus C, Klingman KL, Chege W, Rinehart AR, Rooney JF, Andrew P, Salata RA, Magnus M, Farley JE, Liu A, Frank I, Ho K, Santana J, Stekler JD, McCauley M, Mayer KH. 2017. Phase 2 Study of the Safety and Tolerability of Maraviroc-Containing Regimens to Prevent HIV Infection in Men Who Have Sex With Men (HPTN 069/ACTG A5305). The Journal of Infectious Diseases 215:238–246.

16. Gulick RM, Wilkin TJ, Chen YQ, Landovitz RJ, Amico KR, Young AM, Richardson P, Marzinke MA, Hendrix CW, Eshleman SH, McGowan I, Cottle LM, Andrade A, Marcus C, Klingman KL, Chege W, Rinehart AR, Rooney JF, Andrew P, Salata RA, Siegel M, Manabe YC, Frank I, Ho K, Santana J, Stekler JD, Swaminathan S, McCauley M, Hodder S, Mayer KH. 2017. Safety and Tolerability of Maraviroc-Containing Regimens to Prevent HIV Infection in Women A Phase 2 Randomized Trial. Annals of Internal Medicine 167:384–+.

17. Hill LM, Golin CE, Pack A, Carda-Auten J, Wallace DD, Cherkur S, Farel CE, Rosen EP, Gandhi M, Prince HMA, Kashuba ADM. 2020. Using Real-Time Adherence Feedback to Enhance Communication About Adherence to Antiretroviral Therapy: Patient and Clinician Perspectives. Janac-Journal of the Association of Nurses in Aids Care 31:25–34.

18. Sekabira R, Yuhas K, McGowan I, Brand RM, Marzinke M, Mayer K, Landovitz RJ, Wilkin T, Amico KR, Manabe Y, Frank I, Kekitiinwa AR, Gulick RM, Hendrix CW. 2020. Higher Colon Tissue Infectivity in HIV Seronegative Cisgender Women compared to Cisgender Men on Candidate Oral Antiretroviral (ARV) Pre-Exposure Prophylaxis (PrEP) Regimens in HPTN 069. Journal of the International Aids Society 23:133–134.

19. Rollins DE, Wilkins DG, Krueger GG, Augsburger MP, Mizuno A, O’Neal C, Borges CR, Slawson MH. 2003. The effect of hair color on the incorporation of codeine into human hair. Journal of Analytical Toxicology 27:545–551.

20. Slawson MH, Wilkins DC, Rollins DE. 1998. The incorporation of drugs into hair: Relationship of hair color and melanin concentration to phencyclidine incorporation. Journal of Analytical Toxicology 22:406–413.

21. Gilliland WM, White NR, Yam BH, Mwangi JN, Prince HMA, Weideman AM, Kashuba ADM, Rosen EP. 2020. Influence of hair treatments on detection of antiretrovirals by mass spectrometry imaging. Analyst 145:4540–4550.

22. LeBeau MA, Montgomery MA, Brewer JD. 2011. The role of variations in growth rate and sample collection on interpreting results of segmental analyses of hair. Forensic Science International 210:110–116.

23. Podsadecki TJ, Vrijens BC, Tousset EP, Rode RA, Hanna GJ. 2008. “White coat compliance” limits the reliability of therapeutic drug monitoring in HIV-1-infected patients. HIV Clinical Trials 9:238–246.

24. Asmuth DM, Thompson CG, Chun TW, Ma ZM, Mann S, Sainz T, Serrano-Villar S, Utay NS, Garcia JC, Troia-Cancio P, Pollard RB, Miller CJ, Landay A, Kashuba AD. 2017. Tissue Pharmacologic and Virologic Determinants of Duodenal and Rectal Gastrointestinal-Associated Lymphoid Tissue Immune Reconstitution in HIV-Infected Patients Initiating Antiretroviral Therapy. Journal of Infectious Diseases 216:813–818.

25. Brown KC, Patterson KB, Malone SA, Shaheen NJ, Asher Prince HM, Dumond JB, Spacek MB, Heidt PE, Cohen MS, Kashuba ADM. 2011. Single and Multiple Dose Pharmacokinetics of Maraviroc in Saliva, Semen, and Rectal Tissue of Healthy HIV-Negative Men. The Journal of Infectious Diseases 203:1484–1490.

26. Hendrix CW, Andrade A, Bumpus NN, Kashuba AD, Marzinke MA, Moore A, Anderson PL, Bushman LR, Fuchs EJ, Wiggins I, Radebaugh C, Prince HA, Bakshi RP, Wang R, Richardson P, Shieh E, McKinstry L, Li X, Donnell D, Elharrar V, Mayer KH, Patterson KB. 2016. Dose Frequency Ranging Pharmacokinetic Study of Tenofovir-Emtricitabine After Directly Observed Dosing in Healthy Volunteers to Establish Adherence Benchmarks (HPTN 066). Aids Research and Human Retroviruses 32:32–43.

27. Mayer KH, Yuhas K, Amico KR, Wilkin T, Landovitz RJ, Richardson P, Marzinke MA, Hendrix CW, Eshleman SH, Cottle LM, Marcus C, Chege W, Rinehart AR, Rooney JF, Andrew P, Salata RA, Magnus M, Farley JE, Liu AY, Frank I, Ho K, Santana J, Stekler JD, Chen YQ, McCauley M, Gulick RM, Team HAS. 2022. Sexual behavior and medication adherence in men who have sex with men participating in a pre-exposure prophylaxis study of combinations of Maraviroc, Tenofovir Disoproxil Fumarate and/or Emtricitabine (HPTN 069/ACTG 5305). AIDS and Behavior doi:10.1007/s10461-022-03736-z.

28. Rosen EP, Thompson CG, Bokhart MT, Prince HMA, Sykes C, Muddiman DC, Kashuba ADM. 2016. Analysis of Antiretrovirals in Single Hair Strands for Evaluation of Drug Adherence with Infrared-Matrix-Assisted Laser Desorption Electrospray Ionization Mass Spectrometry Imaging. Analytical Chemistry 88:1336–1344.

29. Mwangi JN, Gilliland WM, Jr., White N, Sykes C, Poliseno A, Knudtson KA, Hightow-Weidman L, Kashuba ADM, Rosen EP. 2022. Mass Spectroscopy Imaging of Hair Strands Captures Short-Term and Long-Term Changes in Emtricitabine Adherence. Antimicrob Agents Chemother 66:e0217621.

30. Gilliland WM, Prince HMA, Poliseno A, Kashuba ADM, Rosen EP. 2019. Infrared Matrix-Assisted Laser Desorption Electrospray Ionization Mass Spectrometry Imaging of Human Hair to Characterize Longitudinal Profiles of the Antiretroviral Maraviroc for Adherence Monitoring. Analytical Chemistry 91:10816–10822.

31. Gerona R, Wen A, Aguilar D, Shum J, Reckers A, Bacchetti P, Gandhi M, Metcalfe J. 2019. Simultaneous analysis of 11 medications for drug resistant TB in small hair samples to quantify adherence and exposure using a validated LC-MS/MS panel. Journal of Chromatography B 1125:121729.

